# A phage weaponizes a satellite recombinase to subvert viral restriction

**DOI:** 10.1101/2022.06.08.495392

**Authors:** Maria HT Nguyen, Zoe Netter, Angus Angermeyer, Kimberley D. Seed

## Abstract

Bacteria can acquire mobile genetic elements (MGEs) to combat infection by viruses (phages). Satellite viruses, including the PLEs (phage-inducible chromosomal island-like elements) in epidemic *Vibrio cholerae*, are MGEs that restrict phage replication to the benefit of their host bacterium. PLEs parasitize the lytic phage ICP1, unleashing multiple mechanisms to restrict phage replication and promote their own spread. In the arms race against PLE, ICP1 uses nucleases, including CRISPR-Cas, to destroy PLE’s genome during infection. However, through an unknown CRISPR-independent mechanism, specific ICP1 isolates subvert restriction by PLE. Here, we discover ICP1-encoded Adi that counteracts PLE by exploiting the PLE’s large serine recombinase (LSR), which normally mobilizes PLE in response to ICP1 infection. Unlike previously characterized ICP1-encoded anti-PLE mechanisms, Adi is not a nuclease itself but instead appears to modulate the activity of the LSR to promote destructive nuclease activity of the LSR’s specific attachment site, *attP*. The PLE LSR, its catalytic activity, and *attP* are additionally sufficient to sensitize a PLE encoding a resistant variant of the recombination module to Adi activity. This work highlights a unique type of adaptation arising from inter-genome conflicts, in which the intended activity of a protein can be weaponized to overcome the antagonizing genome.

## INTRODUCTION

Bacteriophages (phages) are ubiquitous and predicted to outnumber their bacterial prey ten-fold (1). The constant competitive interactions between bacteria and their phage predators have driven the evolution of numerous anti-phage defense mechanisms that target different stages of the phage life cycle (2, 3). To avoid extinction, phages must overcome bacterial strategies that hamper productive phage infection. The rapid acquisition of offensive and defensive mechanisms results in a co-evolutionary arms race that drives microbial evolution (4). Anti-phage mechanisms encoded on mobile genetic elements (MGEs) allow for rapid dissemination of defenses in microbial populations. Some MGEs can protect their bacterial host from phage predation by functioning as virus-like parasites that hijack phage products to promote their own spread (5–8). These subviral parasites, often referred to as satellites, are integrated into attachment sites in their bacterial hosts’ genomes. Upon induction during infection by a specific phage, a subviral parasite is triggered to excise (5, 9, 10), replicate (11, 12), and hijack their inducing phage’s structural components to package their own genome for transduction to naïve neighboring bacterial cells (7, 8, 13, 14). For the bacterial host of this inter-viral conflict, the subviral parasite provides a benefit to the population because the inducing phage is inhibited. In response, the inducing phage may counteract the anti-phage mechanisms inherent to successful hijacking in order to avoid inhibition, resulting in a subcellular arms race.

*Vibrio cholerae*, the causative agent of the diarrheal disease cholera (15, 16), is under constant attack by predatory phages (17–20). To defend against viral attack, strains of epidemic *V. cholerae* have acquired subviral parasites called the <u>p</u>hage-inducible chromosomal island-like <u>e</u>lements (PLEs) that restrict infection by their specific inducing phage, the dominant lytic phage ICP1 (13). Ten distinct PLEs have been identified to date, revealing these elements as a dynamic but persistent part of the MGE repertoire in *V. cholerae* isolated from cholera patients over the last century (20). PLEs share regions of nucleotide conservation but also contain variable modules often shared between two or more PLEs (10, 13). Regardless of variability, all PLEs have a conserved mechanism to block ICP1 infection: PLEs excise from the chromosome upon infection (10, 13), replicate to a high copy number (12), and package their genomes into modified ICP1 capsids to spread via transduction to neighboring *V. cholerae* cells (14) (Figure 1A). PLEs employ multiple mechanisms to interfere with different stages of the ICP1 life cycle that function synergistically to prevent successful phage propagation (14, 19, 21). PLE’s parasitism of ICP1 completely abolishes phage progeny production and therefore enhances *V. cholerae* fitness by protecting the bacterial cell population from further phage attack (12, 13, 19, 21).

**Figure 1.**
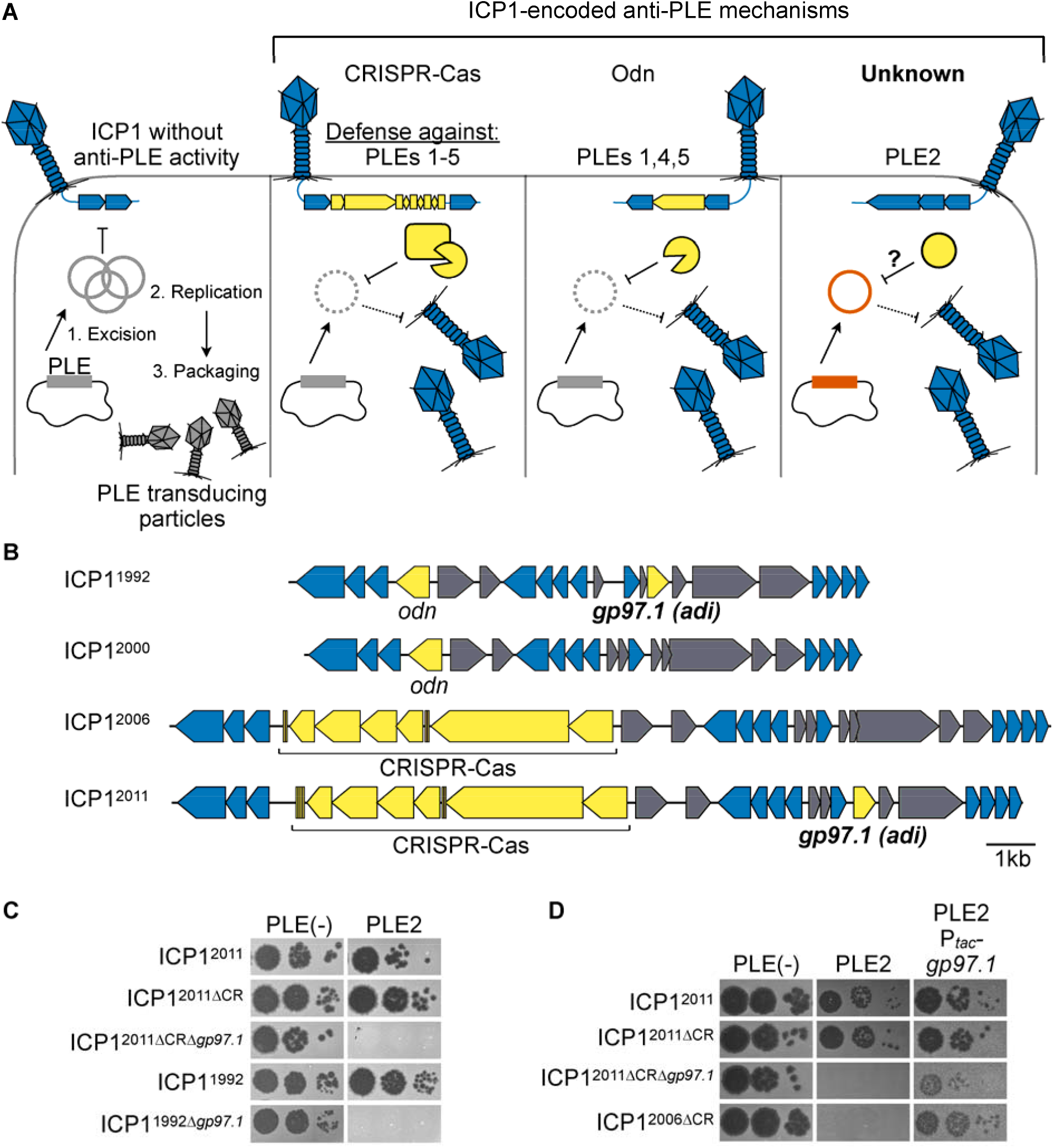
ICP1 isolates encoding *adi* (*gp97*.*1)* overcome PLE2-mediated anti-phage activity. (**A**) Schematic of the molecular warfare between <u>P</u>hage-inducible chromosomal island-<u>L</u>ike <u>E</u>lement (PLE) and the predatory bacteriophage, ICP1. ICP1 infection of PLE(+) *V. cholerae* triggers PLE excision and replication. PLE blocks phage production and packages its genome into transducing particles derived from ICP1 components that are released upon cell lysis. ICP1 overcomes PLE’s anti-phage activity by deploying nucleases: some isolates of ICP1 encode CRISPR-Cas which defends against PLEs 1-5 (5, 19). Alternatively, ICP1 can encode the origin directed nuclease (Odn) that defends against PLEs 1,4 and 5, but not PLE2 or PLE3 (25). A CRISPR(-) derivative of ICP1^2011^ was shown to successfully replicate in the presence of PLE2 specifically (5), suggesting an unknown mechanism for ICP1 to defend against PLE2 (“Unknown” right panel). (**B**) Genomic organization of the regions of the ICP1 phage genome encoding anti-PLE mechanisms in representative isolates. Genes represented by blue arrows are conserved in all ICP1 isolates sequenced to date (26), while accessory genes in yellow have anti-PLE activity. Genes shown in gray are accessory genes that co-vary with the genes involved in countering PLE. (**C**) Ten-fold serial dilutions of various ICP1 isolates encoding *gp97*.*1* and CRISPR (CR) or *odn* and their mutant derivatives spotted onto permissive PLE(-) or PLE2(+) *V. cholerae* (bacterial lawns in gray, zones of killing are shown in black). (**D**) Ten-fold serial dilutions of various ICP1 isolates spotted onto the *V. cholerae* host indicated (bacterial lawns in gray, zones of killing are shown in black). P_*tac*_-*gp97*.*1* is the plasmid expression construct to express Gp97.1 *in trans*. Spot assays were performed in biological triplicate, and a single representative image is shown. For (B, C, and D) the year of isolation for each ICP1 isolate is used as shorthand to differentiate unique isolates (full isolate details provided in Supplementary Table S2).

As an obligate intracellular parasite of *V. cholerae*, ICP1 risks extinction unless it acquires counter-defense(s) against PLE-mediated inhibition. ICP1 isolates, which have been recovered continuously since 1992, share a core genome and differ by accessory components, including those that allow ICP1 to compete against PLE (22, 23). Of these accessory components, all ICP1 isolates encode one of two anti-PLE mechanisms: either a CRISPR-Cas system or a stand-alone nuclease, Odn (18, 22–24) (Figure 1A). CRISPR-Cas uses PLE-derived spacers transcribed from the CRISPR array to guide the effector Cas complex to cleave PLE at various genomic locations, while the nuclease Odn specifically targets the PLE origin of replication. Both of these anti-PLE adaptations directly target the PLE genome, suggesting that the integrity of the PLE genome is integral to combating PLE’s restriction of ICP1. When ICP1 phages are armed with either CRISPR-Cas or Odn, PLE replication and transduction are limited, thereby restoring ICP1 phage production (18, 22). However, the PLE replication module has sufficient allelic variation that allows some PLEs to escape nuclease attack by Odn (12, 18). In this way, it appears that one of ICP1’s counter-defenses against PLEs has directly contributed to PLE diversification. Since CRISPR-Cas and Odn occupy the same genomic locus across all known ICP1 isolates, with a phage encoding either one or the other, ICP1 isolates encoding *odn* would be seemingly incapable of overcoming certain PLEs, like PLE2, which harbors an origin of replication not recognized by Odn (22). Therefore, there may be selection for an additional anti-PLE2 factor that could work synergistically with Odn to broadly inhibit a larger suite of PLEs. Interestingly, an ICP1 isolate with mutations that inactivated its CRISPR-Cas system was still able to replicate and form plaques on *V. cholerae* harboring PLE2, but not other PLEs (13) (Figure 1A). This suggests that ICP1 phage isolates may have additional mechanisms to antagonize certain PLE variants that await discovery.

A conserved aspect of PLE activity is excision from the *V. cholerae* chromosome immediately following ICP1 infection (10, 13). The mobilization of PLE is integral to its life cycle, allowing for its dissemination to naïve *V. cholerae* cells, which then gain the capacity to restrict ICP1 (10, 13). This mobilization is mediated by integrases, which are site-specific recombinases that fall into two protein families based on their catalytic residue: tyrosine or serine (25). Like other MGEs, PLEs rely on integrases to mediate integration and excision in response to specific cues. PLEs encode large serine recombinases (LSRs), which are simple, requiring only the integrase and specific attachment (*att*) sites to carry out integration (25, 26). LSRs possess specific DNA binding activity to recognize attachment sites and catalytic activity to introduce a double-stranded break for precise recombination. Due to the specificity and functionality of LSRs, they have been exploited for applications in genetic engineering (27–34).

PLEs encode one of two types of LSR, referred to simply as the PLE1-type and PLE2-type (20). PLE LSRs differ in their integration sites and cognate recombination directionality factor (RDF), which binds to the LSR to mediate excision (25, 26). So far, only the PLE1-type LSRs have been characterized experimentally (10), though attachment site recognition for PLE2-type LSRs can likely be inferred from the conserved integration site of PLEs encoding this LSR type. For all PLEs, the integrase catalyzes PLE excision out of the chromosome in response to ICP1 infection, resulting in a circularized PLE intermediate. PLE is unable to excise and transduce without the integrase (13), therefore integrases play a vital role in the PLE life cycle by allowing for spread of PLE to neighboring cells (10). As for many PLE genes where divergent gene products carry out a conserved activity, it is unclear what may have contributed to the divergence of the PLE LSRs. The observation that PLEs encode divergent replication modules that can afford an evolutionary advantage upon encountering certain ICP1-encoded counter-defenses suggests that other variations observed between PLEs may be attributable to the ongoing molecular arms race with ICP1.

Motivated by the observation that PLE2 did not restrict a CRISPR-deficient ICP1 isolate, we set out to identify the CRISPR-independent mechanism by which ICP1 can antagonize PLE2 and understand why it does not afford protection against other PLEs tested (13). We found that ICP1 isolates can encode an effector protein whose activity results in degradation of the PLE2 genome. Further, we observed that the anti-PLE activity of this accessory protein is dependent on the PLE2 integrase and the specific target sequence generated by the LSR upon PLE2 mobilization. Unlike known anti-PLE mechanisms used by ICP1, this effector does not appear to be a nuclease itself but rather exploits the satellite-encoded recombinase to execute successful counter-defense. This work broadens our understanding of the innovations resulting from ongoing molecular arms races by revealing that weaponizing the enzymatic activity of other proteins can provide a means of adaptation.

## METHODS

### Strains and culture conditions

A complete list of strains used throughout this study can be found in Supplementary Table S1. *V. cholerae* strains used in this study are derivatives of E7946. Bacteria were grown on LB agar plates with appropriate antibiotic(s) at the following concentrations where appropriate: 75µg/mL kanamycin, 100µg/mL spectinomycin, 2.5µg/mL chloramphenicol (for *V. cholerae* on plates and in 0.5% LB soft agar), 1.25µg/mL chloramphenicol (for *V. cholerae* in liquid), 25µg/mL chloramphenicol (for *Escherichia coli*).

### Phage spot plates

From LB agar plates, 2mL cultures of each *V. cholerae* strain were grown to OD_600_ >1, then back-diluted into fresh media to OD_600_=0.05, and grown to OD_600_=0.2. For *V. cholerae* strains with *adi* under an inducible promoter on a plasmid, cultures were induced with 0.15mM theophylline for 20 minutes at 37ºC with aeration. Then 150µL of mid-log *V. cholerae* was added to 4mL of molten 0.5% LB soft agar (supplemented with 2.5µg/mL chloramphenicol and 0.15mM theophylline), poured onto 100mm x 15mm LB agar plates and allowed to solidify. For *V. cholerae* strains with *int* or *int*^*S12A*^ under an inducible promoter on a plasmid, cultures were induced with 1mM isopropyl-β-D-1-thiogalactopyranoside (IPTG) for 20 minutes. Then 150µL of mid-log *V. cholerae* was added to 4mL of molten 0.5% LB soft agar (supplemented with 2.5µg/mL chloramphenicol and 1mM IPTG), poured onto 100mm x 15mm LB agar plates and allowed to solidify. Once the LB agar plates solidified, 2µL of 10-fold serially diluted phages were spotted onto the plates and incubated for 4 hours at 37ºC. Following incubation, the plates were taken out of the incubator and imaged the next day. Assays were done in triplicate and a single representative image is shown. Replicates have been deposited on Mendeley Data.

### Efficiency of plaquing infection assays

From LB agar plates, 2mL cultures of each *V. cholerae* strain were grown to OD_600_ >1, then back-diluted into fresh media to OD_600_=0.05, and grown to OD_600_=0.2. For *V. cholerae* strains with the empty vector control, or *int* or *int*^*S12A*^ under an inducible promoter on a plasmid, cultures were induced with 1mM IPTG for 20 minutes at 37ºC with aeration. Then 10µL of ten-fold serially diluted phages were added to 50µL of mid-log *V. cholerae* and incubated for 5 minutes to allow for phage attachment to cells. Infected cells were added to 4mL of molten 0.5% LB soft agar (supplemented with 2.5µg/mL chloramphenicol and 1mM IPTG), poured onto 60mm x 15mM petri dishes and allowed to solidify. Once the LB agar plates solidified, the plates were incubated overnight at 37ºC and plaque forming units were counted and compared to the permissive strain to calculate for efficiency of plaquing.

### Generation of mutant strains and constructs

For *V. cholerae* mutants containing chromosomal expression constructs, PCR products were created using splicing by overlap extension (SOE) PCR and introduced by natural transformation as described previously (35). Ectopic expression plasmids are derivatives of pMMB67EH containing a theophylline inducible riboswitch E as described in (10). Gibson assembly was used to assemble plasmid-based expression constructs which were then introduced into *V. cholerae* through conjugation with *Escherichia coli* S17. Mutant derivatives of PLE2 were constructed by inserting a selectable marker containing arms of homology to allow for recombination using the same process detailed above. ICP1 mutants with clean deletions of *adi* were created through CRISPR-Cas gene editing as described previously (36). All constructs were confirmed via Sanger sequencing. The cloned *attP* sequence is provided in Supplementary Table S3.

### Circularization assay

From liquid cultures, *V. cholerae* PLE2(+) or PLE2 mutants were infected with ICP1 phage at an MOI of 1 and incubated with aeration at 37ºC for the time indicated before 100µL of the sample was removed and boiled for 10 minutes. Alternatively, agar stabs of plaques (successful ICP1 infection on *V. cholerae*) were placed in 100µL water and boiled for 10 minutes. 2µL of each boiled sample (from liquid or plaque) was used as template for PCR using primers indicated in Supplementary Table S2. Resulting reactions were run on 2% agarose gel with GelGreen (Biotium) before imaging using an EZ Dock Imager (Bio-Rad). For ectopic expression of Int, *V. cholerae* containing the plasmid expressing Int was induced with 1mM IPTG and 1.5mM theophylline at 37ºC with aeration for 20 minutes at OD_600_=0.2 prior to phage infection. For ectopic expression of *adi* in miniPLE2, *V. cholerae* containing the miniPLE2 construct and plasmid encoding *adi* was induced for 20 minutes beginning at an OD_600_=0.2 prior to being boiled and used as templates for PCR, as described above.

### Toxicity assays

From LB agar plates, 2mL cultures of each *V. cholerae* strain were grown to OD_600_ >1, then back-diluted into fresh media to OD_600_=0.05, grown to OD_600_=0.2 (if slightly greater, normalized to OD_600_=0.2) and induced with 1mM IPTG for 20 minutes at 37ºC with aeration. Induced cultures were plated on LB agar plates supplemented with 1mM IPTG and appropriate antibiotics.

### Transduction assays

A *V. cholerae* PLE2(+) donor with a kanamycin resistance cassette downstream of *orf*27 (as in (13)) was grown to OD_600_=0.3 and infected with ICP1 at a MOI of 2.5. Cultures were incubated with aeration at 37ºC for 5 minutes, centrifuged at 5000 × g for 2 minutes, and washed twice with fresh LB to remove unbound phage. Infected cell pellets were then resuspended in 2mL LB supplemented with 10mM MgSO_4_ and incubated for 30 minutes at 37ºC with aeration. Following incubation, during which time lysis occurred, lysates were treated with chloroform and centrifuged at 4000 × g for 10 minutes to remove debris. 10µL of the transducing lysate was added to 100µL overnight culture of a PLE(−) spectinomycin-resistant (Δ*lacZ*::spec) recipient strain and incubated for 20 minutes at 37ºC with aeration before plating on LB plates supplemented with kanamycin and spectinomycin to select for transductants. Assays were performed in biological triplicate.

### Deep sequencing of phage infected cultures and reads mapping

From LB agar plates, 2mL cultures of each *V. cholerae* strain were grown to OD_600_ >1, then back-diluted into fresh media to OD_600_=0.05, grown to OD_600_=0.3, and infected with ICP1 at an MOI of 1 and then incubated at 37ºC with aeration. 1mL aliquots at t=8 and 16 minutes were taken and immediately pipetted into 1mL ice-cold methanol. Samples were centrifuged at 5000 × g for 3 minutes and total DNA was extracted using the QIAGEN DNeasy Blood and Tissue Kit. Illumina sequencing was performed by the Microbial Genome Sequencing Center (Pittsburgh, PA) at a 200Mbp depth. Then, Illumina sequencing reads for t=8 and t=16 minutes were mapped to the relevant reference files to calculate read coverage and replicates were averaged. Read coverage across all genomic elements (*V. cholerae* chromosomes, ICP1 and PLE2) was reported as percent of total reads normalized by element length. A ratio of averaged coverages for the PLE2 genome was then determined at each nucleotide position between experimental infections with ICP1^ΔCRΔ*adi*^ divided by ICP1^ΔCR^. This ratio calculation was also performed for the plasmid reference with *int*^*S12A*^ divided by WT *int*. These calculations were performed for each timepoint variable and presence/absence of *attP* on the plasmid vector and then log_2_ transformed. The resulting coverage ratios across each reference for every experiment were then scanned using the find_peaks function in SciPy ((37)) with the following settings: height=None, threshold=None, distance=250, prominence=0.2, width=500, wlen=None, rel_height=0.5, plateau_size=None. The five most prominent inverse peaks (i.e. ‘valleys’), which equate to coverage dropouts, were plotted centered on the peak with 250bp on either side.

### Quantification of PLE genome replication by real-time quantitative PCR

From LB agar plates, liquid cultures were grown to OD_600_>1, back-diluted into fresh media to OD_600_=0.05, grown to OD_600_=0.3, infected with ICP1 at an MOI of 2.5 and incubated at 37ºC with aeration. 100µL samples were taken and immediately boiled for 10 minutes at t=0 (uninfected), then at four-minute increments post infection until the final time point at t=20 minutes. Boiled samples were diluted 1:1000 and subsequently used as templates for IQ SYBR (Bio-Rad) qPCR reactions. All conditions were tested in biological triplicate, and each reported data point is the mean of two technical replicates. The fold change was calculated by comparing the Cq value to the standard curve of known concentrations of PLE2 genomic DNA. A single primer set (in Supplementary Table S2) that amplifies a conserved region in all PLEs was used to detect PLE2 replication by qPCR.

### Green Fluorescence Protein (GFP) reporter assays

The *attP* sequence (as in Supplementary Table S2) was engineered between a constitutive promoter and GFP in a neutral locus (*lacZ*) in the *V. cholerae* chromosome. Plasmids containing Int, Int^S12A^ or empty vector control were mated into the GFP-expressing *V. cholerae* strain. From LB agar plates, liquid cultures were grown to OD_600_>1, back-diluted into fresh media to OD_600_=0.05, grown to OD_600_=0.3. Cultures were induced with 1mM IPTG and 1.5mM theophylline for 20 minutes to induce expression of *int* or *int*^*S12A*^. Cells were spun down at 5,000 × g for 3 minutes. Supernatant was removed and washed with 2mL 1X PBS twice. The pellet was resuspended in 2mL 1X PBS and OD_600_ was read and standardized to OD_600_=0.3 across all cultures. 200µL of cultures were transferred to a black 96-well plate and read with an excitation of 485nm and emission of 535nm. Relative fluorescence of *V. cholerae* with *int* or *int*^*S12A*^ was calculated compared to the engineered GFP(+) *V. cholerae* strain containing an empty vector control plasmid. All conditions were tested in biological triplicate, and each reported data point is the mean of two technical replicates.

### Protein purification

*E. coli* BL21 cells containing a 6xHis-SUMO fusion to *int* or *adi*, or 6xHis-SENP2 were grown to OD_600_=0.5 at 37ºC and induced with 0.5mM IPTG and grown for 16 hours at 16ºC. Cells were centrifuged at 4000 × g for 15 minutes and resuspended in 25mL lysis buffer (50mM Tris-HCl pH=7.8 150mM NaCl, 1mM BME, 0.5% Triton-X, 50mM imidazole, Pierce™ protease inhibitor) and sonicated. Cells were centrifuged to remove cell debris. The resulting supernatant was filtered through a 0.2µm cellulose filter (GE Life Sciences). Lysate was applied to an equilibrated HisTrap (Sigma) Ni-sepharose column and washed with 10mL wash Buffer A (50mM Tris-HCl, 150mM NaCl, 50mM imidazole, 1% glycerol, 2mM BME, pH=7.8), followed by 10mL wash Buffer B (50mM Tris-HCl, 2.5M NaCl, 50mM imidazole, 1% glycerol, 2mM BME, pH=7.8), then finally 10mL wash Buffer A. The protein was eluted in 1mL aliquots of imidazole elution buffer (50mM Tris-HCl, 150mM NaCl, 250mM imidazole, 1% glycerol, 2mM BME, pH=7.8). To remove the tag, the eluted protein was dialyzed against wash Buffer A + 5% glycerol overnight in 50K MWCO (Int and IntS12A) and 10K MWCO (Adi) Slide-A-Lyzer™ Dialysis Cassettes (Thermo Scientific™) with 20U/mg of 6xHis-SUMO protease. To remove the SUMO protease and the cleaved tag for Int or Adi, the protein was added to Ni-NTA resin, resulting in the SUMO protease and the cleaved tag binding to the resin and leaving the protein of interest in the unbound buffer. Protein concentration was measured using a BioPhotometer D30 (Eppendorf). For storage, glycerol was added to 20% final concentration and samples were flash frozen for storage at -80ºC.

### Electrophoretic mobility shift assays (EMSAs)

Primers were annealed in water using a thermocycler from 95ºC to 25ºC with an incubation of 1 min per decrease in 1ºC. Primers used as probes are in Supplementary Table S2. 80nM probes were incubated with 500nM purified Int or IntS12A at 30ºC for 20 minutes in 20µL reactions (150mM Tris-HCl, 10mM MgSO_4_, 50mM NaCl, 1mM EDTA, 1mM DTT, pH=7.8). The full reaction volume was then loaded onto an 8% acrylamide 0.5x Tris-borate (TB) gel and ran for 60 minutes at 100V and stained with 0.5x TB buffer with GelGreen (Biotium) before imaging using an EZ Dock Imager (Bio-Rad). Assays were done in triplicate and a single representative image is shown. Replicates have been deposited on Mendeley Data.

### *In vitro* recombination and nuclease assays

Probes were amplified from purified genomic DNA of PLE2(+) *V. cholerae* (*attL* and *attR*), PLE2(+) *V. cholerae* infected with ICP1 phage (*attP*), and PLE(-) *V. cholerae* cells (*attC*). Primers used to amplify probes are available in Supplementary Table S2. PCR products were PCR purified using Monarch® PCR DNA Cleanup Kit (New England BioLabs) and eluted with water. The concentration of probes was measured using a BioPhotometer D30 (Eppendorf). 100ng probes were incubated with purified Int and/or Adi in 20µL reactions (150mM Tris-HCl, 10mM MgSO_4_, 50mM NaCl, 1mM DTT, pH=7.8). Reactions were incubated for 30ºC for 30 minutes followed by 10 minutes at 75ºC to heat inactivate. 10µL of reaction was loaded onto 2% agarose gel with GelGreen (Biotium) and ran for 30 minutes at 100V and visualized using an EZ Dock Imager (Bio-Rad). Assays were done in triplicate and a single representative image is shown. Replicates have been deposited on Mendeley Data.

## Microscopy

*V. cholerae* was grown in LB at 37ºC to OD_600_=0.2, then split to half volumes. Half of each culture was induced with 1mM IPTG, 1.5mM theophylline, and 0.1% arabinose for 20 minutes at 37ºC with aeration. Cultures were pelleted and resuspended in half volume 4% formaldehyde in 0.5x Instant Ocean® supplemented with 20uM Hoechst dye (Thermo Scientific). Samples were incubated in the dark for 10 minutes at room temperature, then pelleted and resuspended in 1/20 volume 0.5x Instant Ocean. 10µL sample volume was mixed with Prolog Gold Antifade mountant (Invitrogen), mounted on a glass slide with a coverslip, and imaged on a Zeiss Axioimager fluorescent microscope with Hamamatsu camera. Experiments were performed in biological triplicate and slides were mounted in technical duplicate and blinded prior to imaging. Exposure times were approximately 30 milliseconds for DIC images and 15 milliseconds for Hoechst-stained DNA fluorescence images. Images were prepared in ImageJ. Assays were done in triplicate and a single representative image is shown. Replicates have been deposited on Mendeley Data.

## RESULTS

### ICP1 encodes adi to overcome PLE2 anti-phage activity

To identify ICP1-encoded gene(s) that allow phage to antagonize PLE2, we began screening for ICP1 phage mutants in the ICP1^2011^ ΔCRISPR background that no longer formed plaques on PLE2(+) *V. cholerae*. Through this screen, we identified *gp97*.*1* (gene product AXY82195.1) as necessary for ICP1^2011^ to form plaques on PLE2(+) *V. cholerae* (Figures 1B and 1C). Gp97.1 is a small protein (147 amino acids) of unknown function unique to ICP1 isolates and has no sequence similarity to proteins of known function. Intriguingly, *gp97*.*1* is encoded just 4.5 kilobase pairs downstream of the locus encoding CRISPR-Cas or *odn* (Figure 1B), suggesting it may provide an additional fitness advantage to phages independent of either of the known anti-PLE mechanisms. Accordingly, *gp97*.*1* was also required for an Odn(+) phage isolate, ICP1^1992^, to form plaques on PLE2(+) *V. cholerae* (Figure 1C). Notably, our initial attempts to complement Δ*gp97*.*1* phage mutants were confounded by the observation that expression of *gp97*.*1* was toxic to PLE2(+) *V. cholerae* (elaborated on further below) (Figure S1). With decreased inducer concentrations (see methods) that allowed the bacterial lawn to form, we found that expression of *gp97*.*1 in trans* restored plaque formation for engineered Δ*gp97*.*1* phage mutants on PLE2(+) *V. cholerae* (Figure 1D). Although *gp97*.*1* is encoded alongside other unique accessory genes (Figure 1B), expression of *gp97*.*1 in trans* was also sufficient to allow the ΔCRISPR derivative of ICP1^2006^ form plaques on PLE2(+) *V. cholerae* (Figure 1D). ICP1^2006^ is an isolate that does not encode *gp97*.*1* or associated flanking genes (Figure 1B), demonstrating that *gp97*.*1* functions independently of co-varying accessory genes. As this gene product is both necessary and sufficient for ICP1 to overcome PLE2, and as informed by subsequent experiments, we named Gp97.1 <u>a</u>ttachment-site <u>d</u>irected inhibitor or Adi.

### Genome replication dynamics following ICP1 infection of PLE2(+) V. cholerae are indicative of Adi-mediated nuclease activity targeting attP

Both known mechanisms that ICP1 uses to counter PLE activity (CRISPR-Cas and Odn) harness nucleases to restrict the robust levels of PLE replication seen during ICP1 infection (12, 24).

Therefore, we evaluated the replication of PLE2 during infection by ICP1 ± *adi*. Moving forward, we performed experiments with ICP1^2011^, which we will refer to as ICP1 for simplicity. We opted to assess replication using deep sequencing of total DNA during an infection time course to provide insight into replication dynamics (12) and reveal potential regions of nuclease cleavage by looking for areas with sudden drops in reads coverage (21). To evaluate total DNA content in infected cells at different stages of infection, we collected samples at relatively early (8 minutes) and late (16 minutes) time points post-infection, consistent with the established kinetics of ICP1 replication and PLE’s response to infection (12, 38). Total DNA from samples at each time point was sequenced and the resulting sequencing reads were mapped to the *V. cholerae*, ICP1, and PLE genomes. Consistent with previous observations measuring replication of a different ICP1 isolate in a permissive PLE(-) *V. cholerae* host (12), ICP1 encoding *adi* infecting a PLE2(+) host led to the robust accumulation of phage DNA late in infection (Figure 2A). As expected for this infection setup wherein PLE2 is antagonized by Adi, robust phage DNA replication coincided with minimal PLE replication. In contrast, during infection by ICP1 Δ*adi*, PLE replication increased ∼7-fold, although replication of PLE2 in these experiments was not as robust as observed previously for PLE1 following infection by a different ICP1 isolate (12). Nevertheless, the increase in PLE replication following infection with ICP1 Δ*adi* coincided with decreased phage replication (Figure 2A), consistent with the established link between PLE replication and inhibition of ICP1 DNA replication (12, 21). The decrease in reads mapping to the PLE genome suggested that PLE transduction would also decrease in the presence of Adi. To test this, we evaluated PLE2 transduction following infection by ICP1 ± *adi*. As expected, PLE transduction efficiency decreased ∼10-fold following infection by *adi*(+) ICP1 compared to the Δ*adi* derivative (Figure S2C). Taken together, the observed replication dynamics and transduction assays are consistent with Adi-mediated interference of PLE replication through nuclease activity directed against the PLE genome itself.

**Figure 2.**
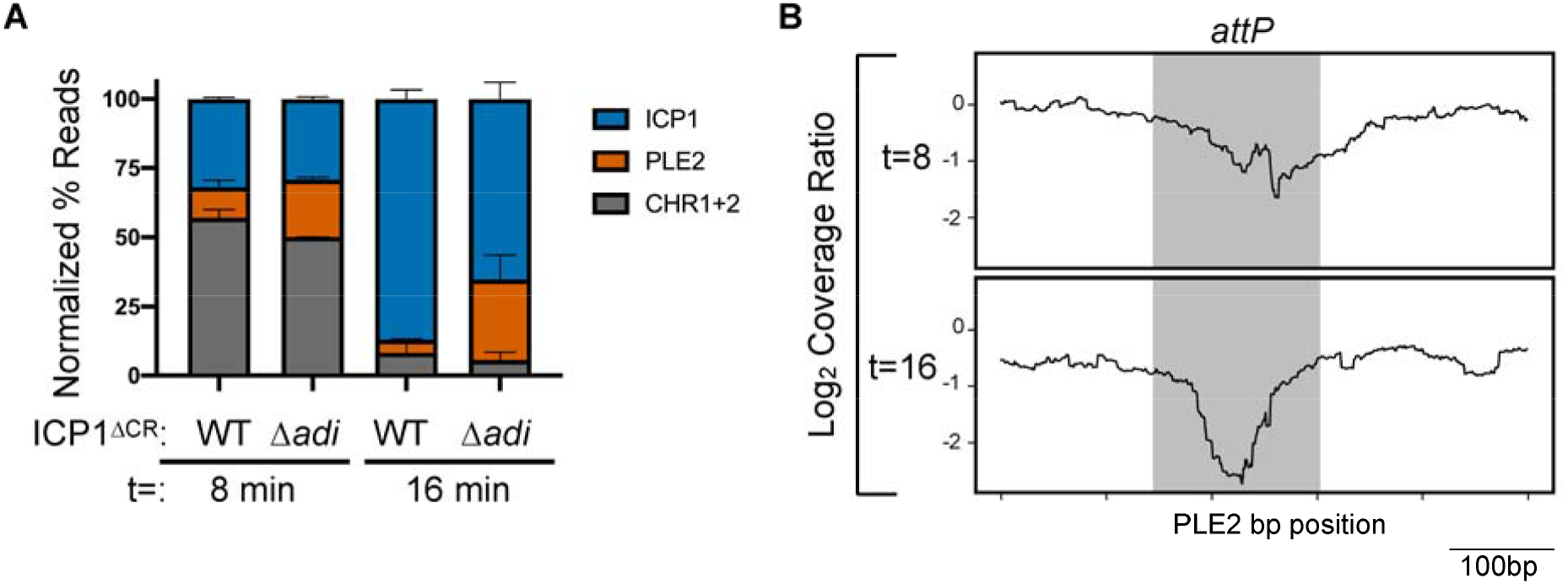
Genome replication dynamics following ICP1 infection of PLE2(+) *V. cholerae* are indicative of Adi-mediated nuclease activity targeting the PLE2 mobilization product, *attP*. (**A**) Percent DNA read abundances normalized to the element’s length for PLE2, both *V. cholerae* chromosomes (CHR1+2), and ICP1 at the time point indicated (in minutes) following infection by ICP1 ΔCRISPR (ΔCR) harboring wild-type Adi (WT) or the Δ*adi* derivative. Bar height is the mean of two biological replicates. (**B**) The log_2_-transformed coverage ratio across the PLE2 genome (relative position shown on the X axis) following infection by Δ*adi* or WT ICP1 over the course of infection. The largest coverage ratio dropout is focused on the 500bp region surrounding *attP* and represents the average of two replicates. The region comprising the circularization junction, *attP*, is highlighted in grey.

Having observed diminished PLE replication during infection with *adi*(+) ICP1, we next determined if there were localized drops in reads coverage in the PLE genome during infection that appeared in an Adi-dependent manner. To generate cleavage maps across the PLE genome, we calculated the coverage ratio of PLE reads obtained following infection with ICP1 Δ*adi* relative to wild-type ICP1 at both 8- and 16-minutes post-infection and analyzed coverage drops, representing a depletion of reads (Figure 2B, Figure S2A and B). Intriguingly, we identified that the reads surrounding the PLE2 circularization junction (*attP* – <u>att</u>achment-site <u>P</u>LE) at both 8- and 16-minutes post-infection showed the largest drop in coverage when ICP1 encoded *adi* (Figure 2B), suggesting nuclease activity targeting this region. Collectively, these data suggest that upon ICP1 infection, PLE2 excises and circularizes to form the *attP* junction, which is then targeted by phage-encoded Adi for degradation, thereby compromising the integrity of the PLE2 genome and favoring productive ICP1 infection.

### Phage-encoded Adi requires the attP target sequence and catalytically active PLE-encoded integrase for anti-PLE2 activity

Following phage infection, the formation of the PLE circularization junction, *attP*, is dependent on the PLE-encoded large serine recombinase (LSR) and recombination directionality factor (RDF) (10, 26). Based on our results, we predicted that Adi mediates PLE interference through nucleolytic cleavage of *attP*; as such, we predicted that PLE2’s LSR would be required for Adi-mediated interference. The PLE1-type LSR (Int) was previously shown to be required for PLE1 excision but was not necessary for PLE to block ICP1 plaque formation (10). To experimentally confirm the requirement of PLE2’s predicted LSR (WP_053027292.1) for PLE excision in response to ICP1 infection, we constructed a PLE2 Δ*int* derivative and challenged it with ICP1. Consistent with the predicted role of the PLE2 LSR, circularization was not detected in the Δ*int* derivative and was restored upon expression of Int *in trans* (Figure S3A). These results were further confirmed by *in vitro* assays using purified PLE2 Int, in which we observed the recombination of *attP* and *attC* sites to form recombinant *attL* and *attR* sites (Figure 3A and Figure S3B), as is characteristic of LSRs (26). As was the case for PLE1 (10), we also found that PLE2 Δ*int* maintained its capacity to block ICP1 plaque formation (Figure 3B).

**Figure 3.**
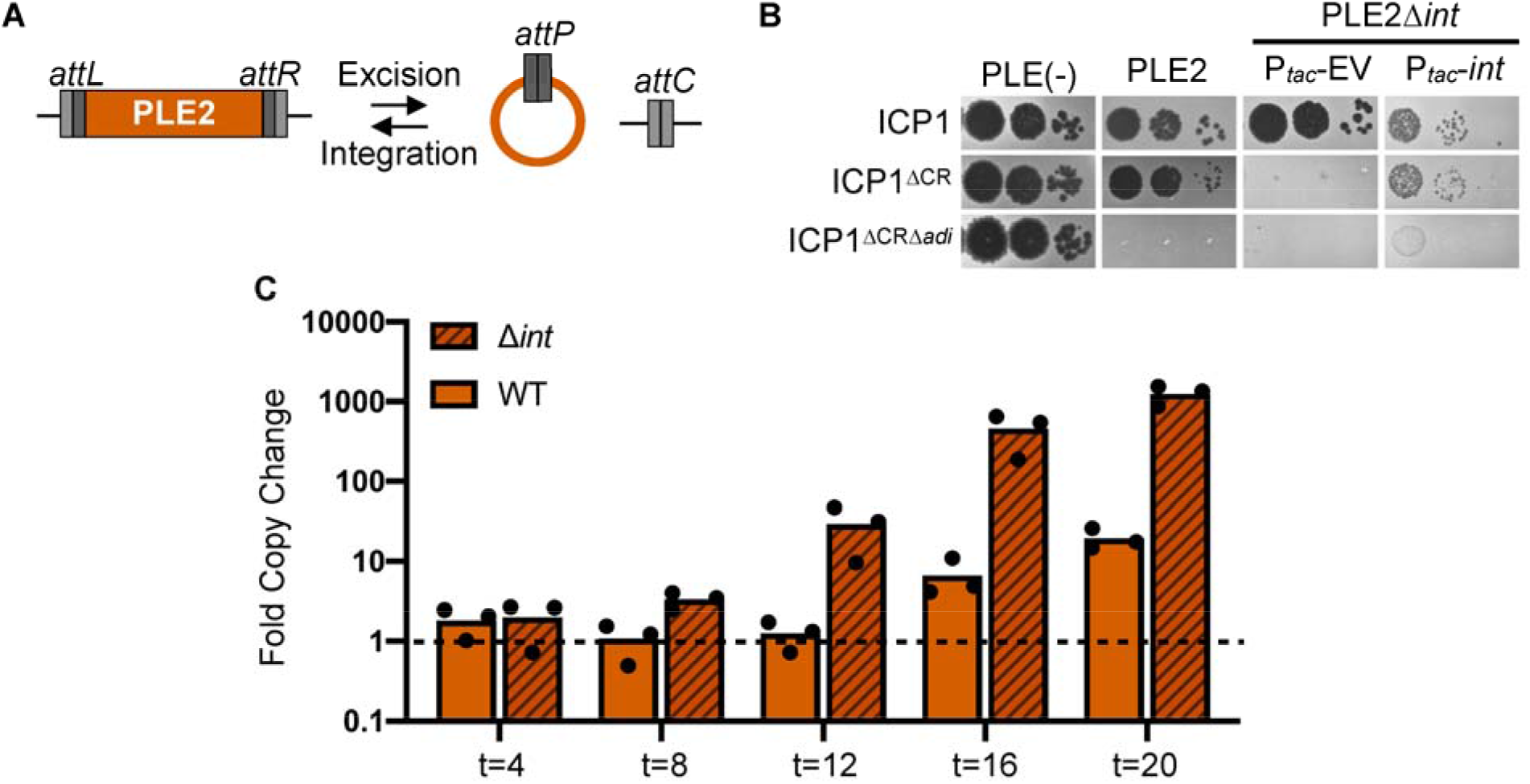
The PLE2 integrase is necessary for Adi(+) ICP1 to antagonize PLE2. (**A**) Schematic of PLE2 integrated into the chromosome with *attL* and *attR* denoting the attachment sites flanking the PLE. Excision of PLE2 upon ICP1 infection results in formation of *attP* and *attC* for the attachment sites on the PLE and the chromosome, respectively. For excision, Int, *att* sites, and recombination directionality factor (RDF) are required. For integration, only the Int and *att* sites are required. (**B**) Ten-fold serial dilutions of Adi(+) ICP1 and the ΔCRISPR (CR), ΔCRΔ*adi* derivatives spotted on PLE(-), PLE2(+) and PLE2Δ*int V. cholerae* containing an empty vector (EV) or plasmid expressing *int* under an inducible promoter. Complemented PLE2Δ*int V. cholerae* were grown and plated with inducer (bacterial lawns in gray, zones of killing are shown in black). (**C**) Replication efficiency of wild-type (WT) PLE2 and the Δ*int* derivative in *V. cholerae* host strains calculated as the fold change in PLE2 DNA copy at the time indicated (in minutes) following infection with Adi(+) ICP1^ΔCR^ as assessed by qPCR.

In the absence of Int, PLE2 does not excise to form the circularization junction *attP* (Figure S3A). Following these observations, we predicted that Adi would require Int to mediate excision of PLE2 to form the target sequence, *attP*, allowing for ICP1 plaque formation. In support of this, we observed that ICP1 encoding *adi* could no longer form plaques on the PLE2 Δ*int* derivative (Figure 3B). ICP1’s plaquing phenotype was restored by expressing Int *in trans*, revealing that PLE2 excision sensitizes it to Adi-mediated anti-PLE activity (Figure 3B).

Replication of PLE1 following phage infection was previously shown to occur independently of PLE excision (10), hence we next evaluated if Int negatively impacted PLE2 replication following infection by ICP1 expressing Adi. Consistent with the role of Int in sensitizing PLE to Adi-mediated interference, we observed that upon Adi(+) ICP1 infection, replication of wild-type PLE2 was impaired ∼100-fold relative to PLE2 Δ*int* (Figure 3C). This observation demonstrates that dependence on the PLE2 LSR is a critical facet of Adi-mediated anti-PLE activity and explains the observed specificity of CRISPR-deficient Adi(+) ICP1, which overcomes PLE2 but was unable to overcome PLEs encoding the PLE1-type LSR (13).

Having observed a drop in reads coverage consistent with cleavage of *attP* in the presence of Adi (Figure 2B), we next sought to test the hypothesis that *attP* alone was sufficient for Adi-mediated targeting of PLE. To do this, we engineered the target sequence *attP* directly into PLE2 Δ*int* (Figure 4A), reasoning that the integrase would now be dispensable for ICP1 plaque formation.

**Figure 4.**
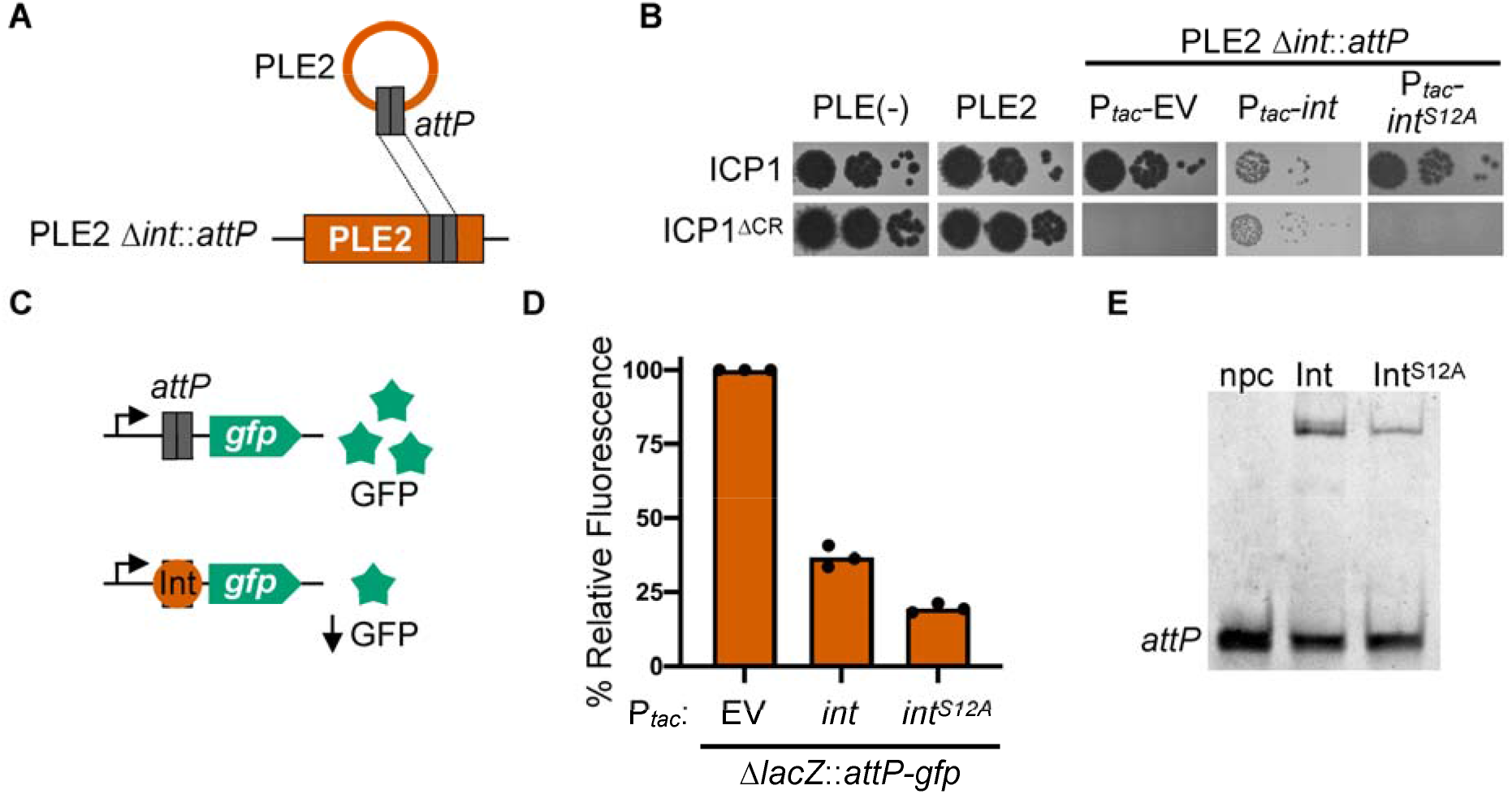
PLE2 *attP* and catalytic activity of the PLE integrase are required for Adi(+) ICP1 to antagonize PLE2. (**A**) Schematic of PLE2 Δ*int::attP* construct with *attP* cloned into PLE2. (**B**) Ten-fold serial dilutions of Adi(+) ICP1 and the ΔCRISPR (ΔCR) derivative spotted on PLE(-), PLE2(+) or PLE2Δ*int*::*attP* containing an empty vector (EV) or plasmid expressing *int* or *int*^*S12A*^ under an inducible promoter grown and plated with inducer (bacterial lawns in gray, zones of killing are shown in black). (**C**) Schematic of the *in vivo* green fluorescence protein (GFP) reporter assay. Steric hindrance by proteins that bind DNA in the region between the constitutive promoter and *gfp* result in lower levels of GFP expression. (**D**) *V. cholerae* cells containing an empty vector (EV) or plasmid expressing *int* or *int*^*S12A*^ under an inducible promoter were grown with inducer. Relative fluorescence of *int* and *int*^*S12A*^ to the empty vector control is shown. Bar height is the mean of three biological replicates and each dot is a measurement from an independent assay. The empty vector control (set at 100%) is shown for reference. (**E**) Electrophoretic mobility shift assay (EMSA) using a probe harboring *attP* incubated without protein (npc), or with purified Int or Int^S12A^.

Surprisingly, we observed that Adi(+) ICP1 was unable to plaque on the *attP*(+) strain, indicating that *attP* was not sufficient to direct Adi activity (Figure 4B). Furthermore, Adi does not possess domains predicted to confer DNA binding or nuclease activity, consistent with *in vitro* assays in which we detected neither activity for purified Adi (Figure S3C and S3D).

Therefore, we next considered the possibility that the requirement of PLE2 Int for Adi-mediated anti-PLE activity (Figure 3B) extended beyond its role in generating the *attP* junction. Notably, ICP1 plaque formation was restored on the *attP*(+) PLE2 Δ*int* host with the complementation of Int *in trans* (Figure 4B), suggesting that the integrase protein itself is required, and ICP1’s inability to plaque on PLE2 Δ*int* is not solely due to the lack of the *attP* target sequence.

Previous work has shown that the catalytic activity of LSRs is decoupled from DNA binding activity (26, 39–41). Therefore, we considered the possibility that Adi may exploit the DNA binding activity of Int to localize to *attP* and target it for degradation. To test this, we engineered a mutation in the predicted catalytic serine residue of PLE2 Int (S12A). We verified that Int^S12A^ was catalytically inactive using an *in vitro* recombination assay where, as expected, purified Int^S12A^ did not recombine *attP* and *attC* (Figure S3D). To verify that Int^S12A^ retained its DNA binding activity, we performed *in vivo* and *in vitro* assays. To assess DNA-binding *in vivo*, we cloned *attP* in between a constitutive promoter and green fluorescence protein (GFP) in the *V. cholerae* chromosome, reasoning that expression of a protein that binds *attP* would result in steric hindrance and decreased fluorescence (Figure 4C). We found that expression of both Int and Int^S12A^ resulted in decreased fluorescence compared to empty vector control, consistent with DNA binding activity for both proteins (Figure 4D). To confirm our *in vivo* assay, we assessed if purified Int and Int^S12A^ could bind *attP* directly using an electrophoretic mobility assay (EMSA) and found that both purified proteins demonstrated binding to *attP in vitro* (Figure 4E). With the catalytically inactive Int^S12A^ we were then able to test whether Adi relies solely on Int’s DNA binding activity, reasoning that complementation of PLE2 Δ*int::attP* with the Int^S12A^ mutant should restore plaque formation if Adi only required Int to localize to *attP*. However, we observed that *adi*(+) ICP1 was still unable to form plaques even with the target sequence *attP* and the DNA binding activity of Int^S12A^ (Figure 4B). Together, these results indicate that ICP1-encoded Adi requires the catalytic activity of the PLE2 LSR and *attP* to exert anti-PLE activity.

Recombination directionality factors (RDFs) are the only group of proteins described to bind directly to LSRs, resulting in a conformational change in the LSR to favor the excision reaction and the formation of the circularized junction, *attP* (26, 42, 43). The RDF for the PLE1-type LSR has been identified as ICP1-encoded PexA (10); however, the RDF for PLE2-type LSRs has not been identified. Our results up to this point indicate that Adi may interact directly with PLE2 Int to induce a conformational change and modulate its activity, similar to the role of an RDF. To address if Adi functions as an RDF for PLE2, we first constructed a minimal platform to assess PLE2 excision, referred to as ‘miniPLE2’, as was previously published for PLE1 (10). MiniPLE2 encodes Int under the control of its endogenous promoter but no other PLE2 genes and is integrated into the same attachment site as full-length PLE2. Upon ICP1 infection, miniPLE2 excised and circularized in the absence of other PLE products, demonstrating that no other PLE2 genes are required for PLE2 excision (Figure S4A). To evaluate if Adi can function as the RDF for PLE2 Int, we expressed *adi* in *V. cholerae* carrying miniPLE2 and found Adi was not sufficient for excision and circularization (Figure S4A). In addition, we observed PLE2 circularization following infection with ICP1 Δ*adi* (Figure S4B), supporting our conclusion that Adi is indeed not the RDF. Having identified at least some of the necessary components for Int/Adi mediated anti-PLE activity, we were motivated to recapitulate the predicted *att*-directed nuclease activity of purified Int and Adi *in vitro*. However, we observed that while Int maintained its recombinase activity *in vitro* in the presence of Adi, there was no obvious aberrant nuclease activity when the purified proteins were combined (Figure S5). Adi also did not inhibit the integration reaction as an RDF would (41, 44, 45), further confirming it is not the RDF. The inability to reconstitute destructive nuclease activity *in vitro* could be due to a missing factor (e.g., other proteins or cofactors in the nuclease buffer). Although we predict that Adi may share features with the PLE2 RDF, namely its ability to interact with and perhaps modulate Int-directed DNA manipulation, these data demonstrate that Adi is not functionally equivalent to the RDF but instead represents a novel factor also capable of altering LSR activity.

### PLE2 Int and attP are sufficient to sensitize PLE1 to Adi-mediated anti-PLE activity

Previously, PLE1 was shown to block plaque formation by *adi*(+) ICP1 (13). This observation aligns with the differences between the mobility module (comprised of Int and cognate attachment-sites) of PLE1 and PLE2. Therefore, we hypothesized that expression of PLE2 Int *in trans* would sensitize PLE1 to *adi*(+) ICP1, but only if the putative target site, *attP*^PLE2^, was cloned into PLE1. To test this, we cloned *attP* directly into PLE1 (Figure 5A) and assessed the sensitivity of this strain to phage infection in the presence and absence of ectopic expression of Int. The *attP*(+) PLE1 strain retained the ability to restrict *adi*(+) ICP1, confirming that *attP* is not sufficient for Adi-mediated anti-PLE activity (Figure 5B). Likewise, PLE1 lacking *attP* retained the capacity to block plaque formation by this phage with Int expressed *in trans* (Figure 5B). In contrast, the PLE1 strain bearing the *attP* sequence was no longer inhibitory to ICP1 with PLE2 Int expressed *in trans* (Figure 5B), confirming that both *attP* and the PLE2 integrase are required to mediate sensitivity to *adi*(+) ICP1. Further, *attP* was not sufficient to sensitize PLE1 to ICP1 antagonism even in the presence of Int if *attP* is cloned outside of PLE in the *V. cholerae* chromosome (Figure 5C and Figure 5D). The requirement of *attP* to be encoded *in cis* (within PLE) is consistent with *attP* serving as the target sequence for Adi-mediated Int-dependent anti-PLE activity.

**Figure 5.**
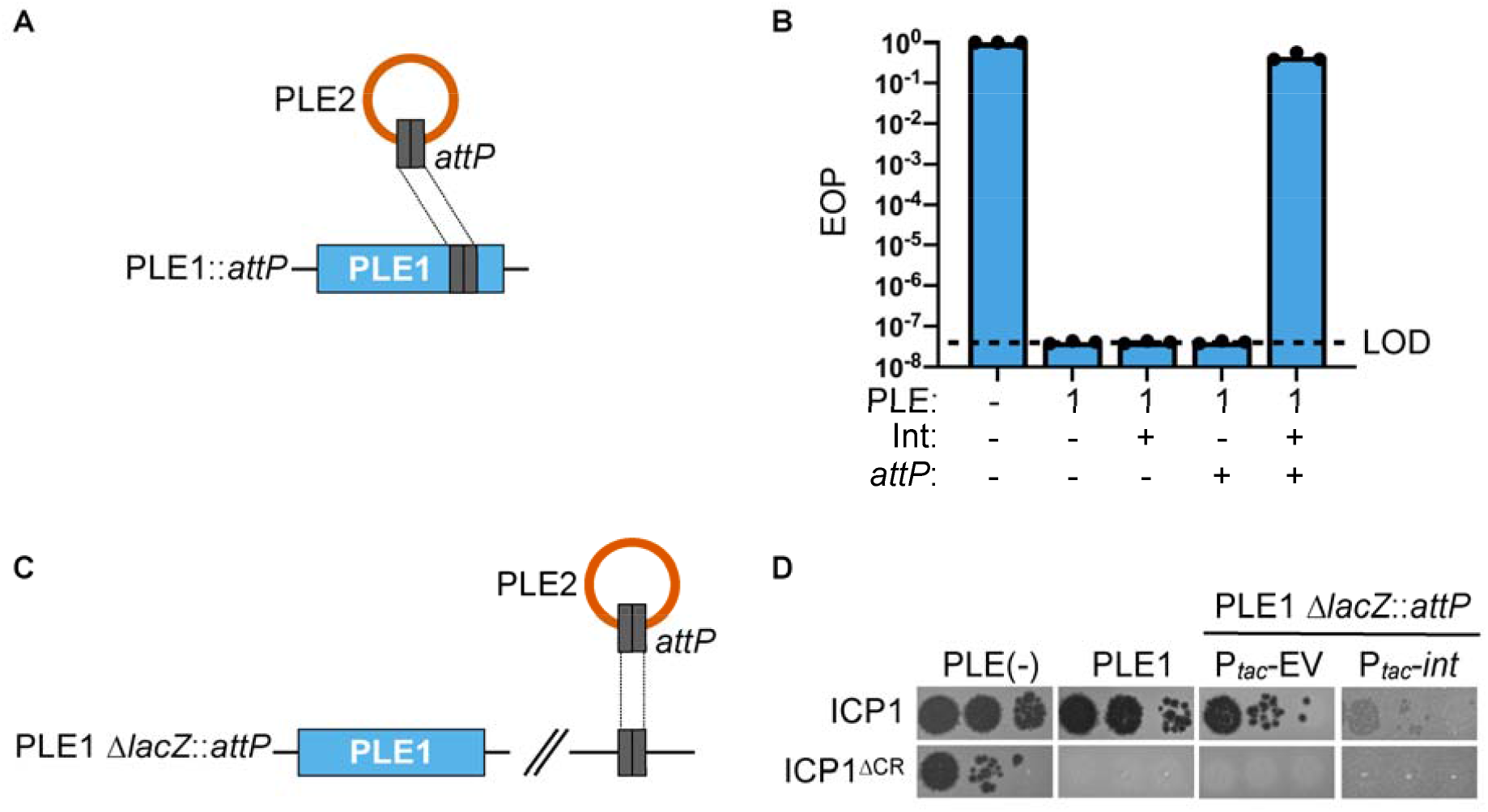
PLE2 Int and *attP* are required to sensitize PLE1 to ICP1 phage infection. (**A**) Schematic showing relevant construct design of PLE1::*attP* wherein PLE2 *attP* was engineered into PLE1. (**B**) Efficiency of plaquing (EOP) of Adi(+) ICP1 ΔCRISPR on *V. cholerae* with the PLE indicated. The presence of an induced plasmid expressing Int from PLE2 is indicated by +/-Int, and the presence of PLE2 *attP* in PLE1 (as in panel A) is indicated by +/-*attP*. EOPs were calculated relative to a permissive PLE(-) host (shown at left for reference). Bar height is the mean of three biological replicates and each dot is a measurement from an independent assay. The dashed line indicates the limit of detection (LOD) for this assay. (**C**) Schematic showing relevant construct design for PLE2 *attP* cloned in a neutral location (*lacZ*) in the *V. cholerae* chromosome. (**D**) Ten-fold serial dilutions of Adi(+) ICP1 and ICP1 ΔCRISPR (ΔCR) spotted on PLE(-), PLE1(+), PLE1(+) Δ*lacZ::attP V. cholerae* containing an empty vector (EV) or *int* under an inducible promoter grown with inducer (bacterial lawns in gray, zones of killing are shown in black).

### Co-expression of Int and Adi results in attachment-site DNA degradation

Having established that Int, Adi, and the target sequence *attP* are sufficient for anti-PLE activity that results in impaired replication of the PLE genome, we wanted to directly assess the fate of DNA in the presence of both Int and Adi. Since we could not recapitulate DNA degradation *in vitro*, we turned to experiments in *V. cholerae* in the absence of phage infection. This was partly motivated by the observation that co-expression of *int*, but not the catalytically inactive mutant *int*^*S12A*^, and *adi* was toxic to *V. cholerae* (Figure 6A). We anticipated this toxicity was due to nuclease activity driven by Int and Adi. To assess this possibility, we performed light and fluorescence microscopy on cells after a short induction of *int, adi*, or both. Congruent with the toxicity assay results, we observed that expression of either *int* or *adi* alone resulted in normal DNA staining, while co-expression of *int* and *adi* resulted in a dramatic loss of DNA signal by Hoechst staining (Figure 6B). By light microscopy all induced cells appeared physiologically normal, suggesting that the specific observed loss of DNA signal during Int and Adi co-expression was not due to apparent cell lysis, membrane damage, or cell division defects. Interestingly, the strain used in these assays is PLE(-) and therefore does not contain *attP*, but it has *attC*, the chromosomal integration site also recognized by Int, suggesting that Int and Adi will destroy any DNA that Int has inherent specificity for.

**Figure 6.**
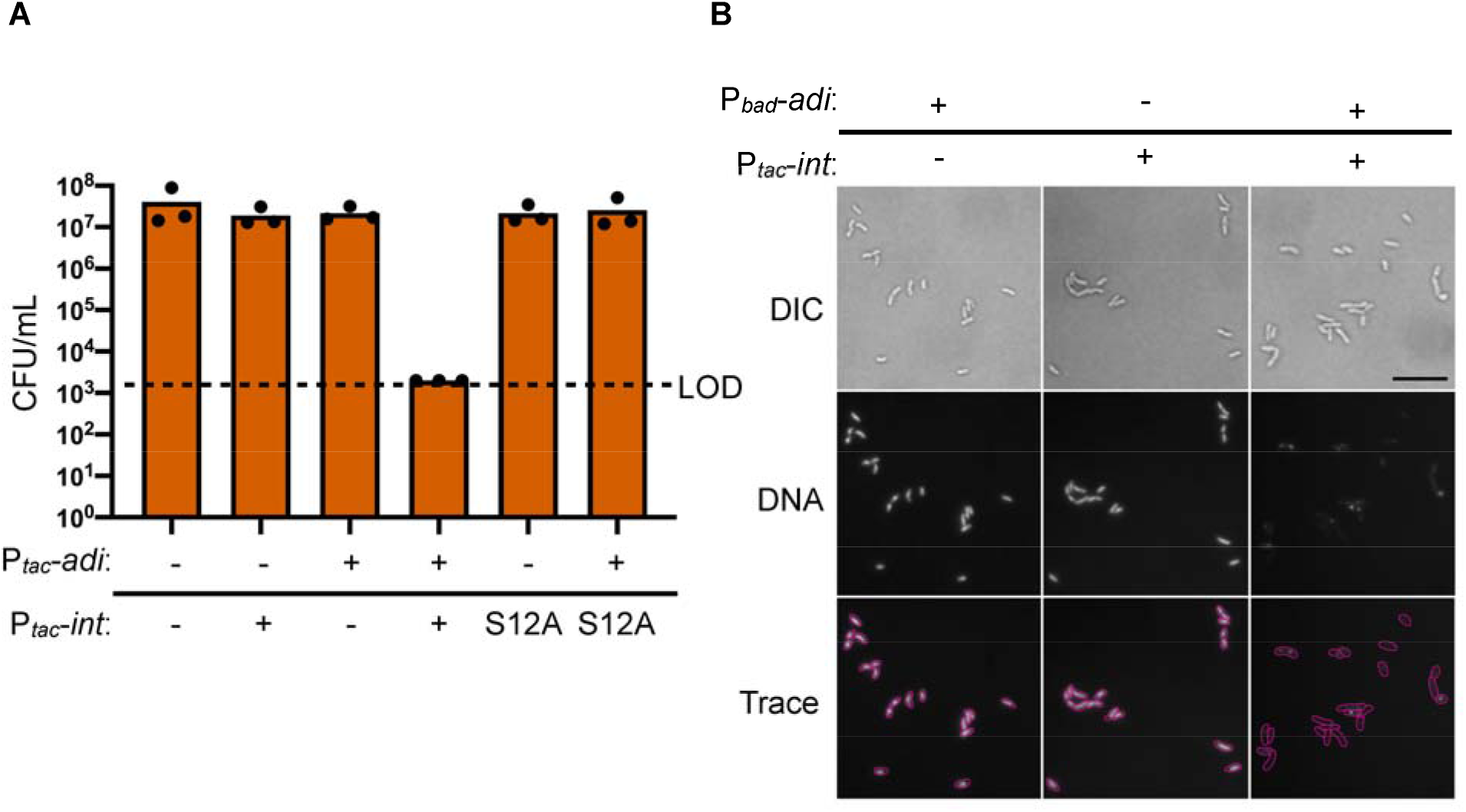
Co-expression of *int* and *adi* results in destructive nuclease activity in *V. cholerae*. (**A**) Cell viability as measured by colony forming units (CFU) per mL of *V. cholerae* expressing *int, int*^*S12A*^, or an empty expression construct (-) in the chromosome with an empty vector plasmid (-) or plasmid expressing *adi* as indicated. Dashed line indicates the limit of detection (LOD). Bar height is the mean of three biological replicates and each dot is a measurement from an independent assay. (**B**) Representative images of differential interference contrast (DIC) light microscopy and fluorescence microscopy of *V. cholerae* cells induced to express either *int, adi*, or both for 20 minutes followed by aldehyde fixation and staining with Hoechst DNA stain. Cells were imaged at 100x magnification with white light (DIC) or DAPI filter (DNA). Scale bar represents 10 microns. Traces were generated manually from matched overlayed DIC and DNA images. Slides were prepared and imaged from three independent biological replicates in technical duplicate and blinded prior to imaging.

The microscopy data and cell survival assays are suggestive of destructive nuclease activity upon co-expression of *int* and *adi* in *V. cholerae* in the absence of other PLE and phage products. However, we reasoned that such nuclease activity must be directed to not be self-destructive for ICP1. Therefore, we were curious whether co-expression of *int* and *adi* was sufficient to mediate DNA degradation specifically of the *attP* sequence, recapitulating the depletion of PLE2’s *attP* observed during phage infection (Figure 2B). To assess this, we co-expressed *int* or the *int*^*S12A*^ mutant with *adi* in *V. cholerae* containing an empty vector control or a plasmid containing the *attP* target sequence and performed deep sequencing of the total DNA before and after induction. The resulting sequencing reads were mapped to the relevant plasmid sequence to look for evidence of plasmid degradation. Strikingly, the proportion of total reads mapping to the *attP*(+) plasmid decreased ∼10-fold only when catalytically active *int* was co-expressed with *adi* (Figure 7A). Importantly, we did not observe depletion of the empty vector plasmid upon co-expression of *int* and *adi*, indicating that the *attP* sequence is required for targeting. We compared the coverages between *V. cholerae* expressing the catalytically active *int* or its mutant derivative *int*^*S12A*^ and scanned for dropouts in the coverage ratio as was performed for PLE2 previously. As expected, we observed that the largest drop in the coverage ratio was localized at the target sequence, *attP* (Figure 7B). Further, we observed no drop in coverage ratio when comparing the induced empty vector plasmid controls, suggesting that nuclease activity specifically targets the *attP*-site (Figure S6). Altogether, these data suggest that Int and Adi are sufficient for *attP* targeting in the absence of phage infection and that no other PLE or ICP1 products are required for the depletion of *attP*.

**Figure 7.**
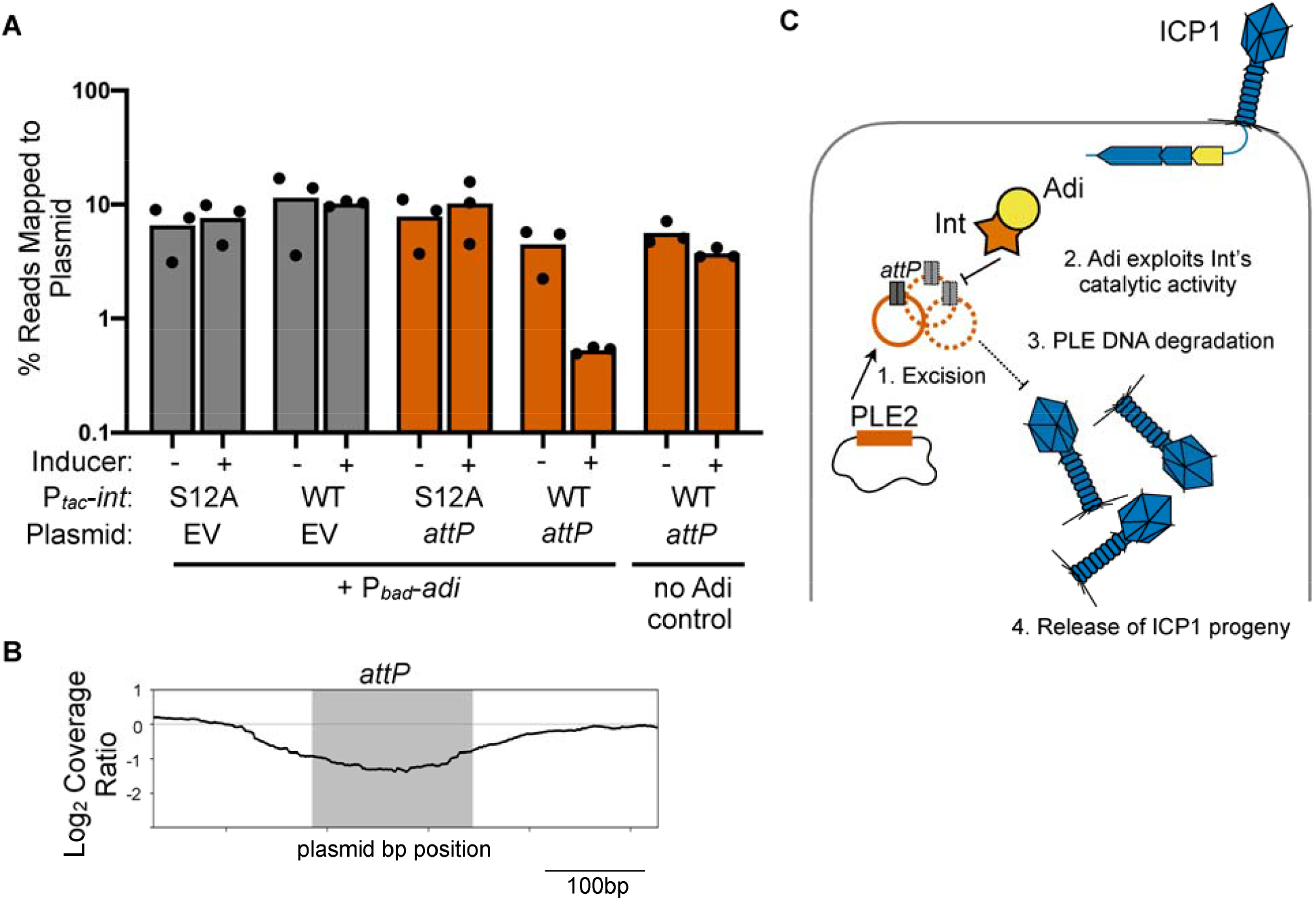
Co-expression of *int* and *adi* targets plasmids in an *attP* dependent manner. (**A**) Percent reads abundance mapped to the plasmid indicated prior to induction (-) and 20 minutes post-induction (+) in *V. cholerae* strains induced to express either *int or int*^*S12A*^ and *adi*. Strains had either an empty vector control (EV) or plasmid containing the PLE2 *attP* target sequence. Bar height is the mean of three biological replicates and each dot is a measurement from an independent assay. (**B**) Log_2_-transformed sequence coverage ratio across plasmid containing *attP* (relative position shown on the X axis) following induction of *int* and *adi* by showing the most prominent coverage drop detected compared to induction of *int*^*S12A*^ and *adi* 20 minutes post-induction. The largest coverage ratio dropout is focused on the 500bp region surrounding *attP* and represents the average of three replicates. The region comprising the circularization junction, *attP*, is highlighted in grey. (**C**) A model representing a possible mechanism of *adi*(+) ICP1 infection exploiting Int-mediated PLE2 mobilization. Upon ICP1 infection, Int excises PLE2 from the chromosome, Adi exploits Int’s catalytic activity resulting in PLE DNA degradation through targeting of *attP*. ICP1 can then successfully replicate and release phage progeny following cell lysis.

Overall, our data support a model in which Adi(+) ICP1 infects a PLE2(+) *V. cholerae* host, resulting in Int-mediated PLE2 excision and the formation of the PLE mobilization product, *attP*. Concomitantly, the anti-PLE factor Adi alters Int activity such that *attP* serves as the target site for its destructive nuclease activity, resulting in loss of integrity of the PLE genome and restored production of ICP1 progeny phage (Figure 7C).

## DISCUSSION

Here, we uncover a novel anti-PLE mechanism present in some ICP1 isolates that appears to function as a counter-defense mechanism by exploiting PLE’s mobility. Thus far, known ICP1-encoded anti-PLE counter-defenses rely on nuclease activity to target the PLE genome: CRISPR-Cas provides anti-PLE defense against a broad range of PLEs (13, 18, 24), while Odn targets the origin of replication of PLEs 1,3,4, and 5 (22). Because Odn does not target all PLEs, Adi serves to expand a given phage’s capacity to defend against all PLEs. This is exemplified by ICP1^1992^, which is armed with both Odn and Adi, allowing it to target multiple PLEs: Odn targets the origin of replication of PLEs 1,3,4,5; while Adi restricts PLE2 by targeting the PLE2 specific *attP* site. Intriguingly, many ICP1 isolates encoding CRISPR-Cas also encode Adi (23), signifying that there is a fitness benefit for ICP1 to maintain Adi, the anti-PLE2 effector, even in the presence of a broader anti-PLE defense system. Although CRISPR-Cas restores ICP1 plaque formation on a PLE2(+) host (13), this defense alone may not be enough to completely abolish PLE replication and transduction, as was observed for PLE1 (18, 24). By encoding CRISPR-Cas and Adi, the combined action of two anti-PLE counter-defense mechanisms could help to compensate for differences in interference seen with different CRISPR spacers (24). Further, encoding two anti-PLE systems would require multiple counter-adaptations by PLE to escape, ensuring ICP1 maintains the upper hand. Alternatively, the nuclease activity stimulated by Adi and Int helps to generate substrates for adaptation of the phage CRISPR system, analogous to what occurs in bacteria with CRISPR and restriction-modification systems (46). Previous analyses show that proteins with similar functions (e.g., anti-defense systems) cluster together in specific genomic locations or hotspots (22, 47–49). Finding *adi* near known anti-PLE genes provides further evidence for a guilt-by-association model of counter-defense gene organization in ICP1. Now that the *adi* locus has been functionally identified, genes in the same genomic context in Adi(-) ICP1 isolates are compelling candidates to explore as possible additional counter-defense mechanisms.

It is interesting as to why PLEs would encode PLE2-type integrases when doing so would sensitize PLE to Adi. Previous work has shown that PLE1-type integrases exploit ICP1-encoded PexA as the RDF to favor the excision of PLE1 from the chromosome (10). However, PexA is not essential and can be mutated in ICP1 (10). While PLE2’s RDF has not yet been identified, it is possible that PLE2-type integrases evolved to exploit an essential ICP1 gene product as their RDF. Therefore, if PLE2-type integrases evolved to exploit an essential gene, then in a tit-for-tat model, perhaps ICP1 evolved to exploit this integrase to PLE’s demise. While the evolutionary path that led to the divergence of mobility modules in PLEs can only be speculated, it is clear that the modular diversity observed across PLEs can provide a means to escape the molecular warfare with their inducing phage while maintaining their fundamental life cycle. Innovations arising from the conflict between these antagonizing genomes continue to inform the investigation of the so-called microbial dark matter, genes and systems whose functions cannot be bioinformatically inferred but become apparent when examined in the context of inter-microbial conflict.

A question remains as to the specific mechanism of Adi activity that renders PLE2 unable to restrict ICP1. Although it is still unclear how Adi modulates Int activity, consideration of how LSRs function may provide insight into the possible mechanism employed by Int/Adi targeting. A hallmark of the LSR mechanism is its concerted action in double-stranded cleavage (26, 39, 45). During excision, LSRs bind to their DNA substrates and form a synaptic complex leading to activation of the catalytic serine residue and cleavage of the DNA substrates at the crossover dinucleotide (26, 39, 45). An intermediate of this recombination is the formation of dinucleotide 3’ overhangs (26, 39, 45). One of the possible mechanisms by which Adi could act to damage the PLE2 genome is by preventing the integrase from re-ligating the dsDNA break while the integrase is mediating recombination, thereby resulting in free 3’ hydroxy groups on DNA ends that nucleases could target. While this proposed mechanism suggests a direct interaction between Int and Adi, we did not see evidence of an interaction using pulldown assays or bacterial adenylate cyclase two-hybrid assays. Our attempt to recapitulate the protein-protein interaction may be confounded by the transience of the interaction or the lack of RDF (or an additional factor) that could stabilize the interaction between Int and Adi. Notably, however, expression of Int and Adi in *V. cholerae* is sufficient for *attP* targeting, suggesting that nuclease activity targeting the PLE does not require other PLE or ICP1-encoded factors (including the putative RDF) but does not rule out requirements for other *V. cholerae* host-encoded factors. Lastly, another way in which Int activity can be modulated is through modification of the protein itself. However, we are not aware of any LSRs that can undergo modification, except during recombination in which the LSR is covalently bound to the 5’ DNA ends through a phosphoseryl linkage (26, 39–41). Thus, further study of the mechanism by which Adi exploits Int activity to target PLE2 may reveal more widespread modifications to LSR activity than are currently appreciated.

To our knowledge, RDFs are the only known proteins that interact with LSRs to modulate their activity. In the context of phage infection, Int binds to the RDF resulting in PLE excision. At the molecular level, RDF-Int binding alters the activity of the LSR, modulating Int’s activity to favor excision resulting in the recombination between *attL* and *attR* (44, 50, 51), and inhibiting *attP* and *attC* recombination (integration) (44). Because of the simple requirements and specificity of directional recombination, LSRs are used as tools for genetic engineering (27–34). The discovery of Adi and the data presented here suggest that other factors could modulate the activity of LSRs in unexpected ways (i.e., to have destructive nuclease activity) and implicates an additional barrier in the development of LSRs as tools for genetic engineering.

Our model of Adi activity raises the possibility that other factors, including known defenses, could be exploited in inter-genome conflicts. Nucleases are particularly pervasive effectors mediating conflicts between hosts, mobile elements, and viruses (52). Such nucleases could be manipulated to the detriment of the organism deploying them, in a similar manner to how Adi exploits Int for anti-PLE activity. For instance, perhaps some anti-CRISPRs bind Cas proteins to modulate their activity in such a way that instead of cleaving the antagonizing genome, Cas targets its host genome for degradation instead. Likewise, restriction-modification systems could be exploited in such a way that the self-genome usually protected from nuclease destruction becomes targeted. Moreover, such proteins that alter the activity of other enzymes could be challenging to predict, perhaps explaining why such mechanisms have yet to be discovered. Still, encoding small proteins like Adi that weaponize existing proteins or even large protein complexes may allow additional pathways to survival for mobile elements like phages that have constrained genome sizes.

## DATA AVAILABILITY

The sequencing data from phage (± *adi* (*gp97*.*1))* infected PLE2(+) *V. cholerae* and plasmid depletion assays generated in this study have been deposited in the Sequence Read Archive database under BioProject accession code PRJNA839547. Replicates have been deposited to Mendeley Data, during review these data can be accessed at the following link https://data.mendeley.com/v1/datasets/2nzzpp69g9/draft?a=f6cb66f3-efb7-4e78-a46f-cb9b3db1362b.

## ACKNOWLEDGEMENTS

We thank members of the Seed lab, including former lab member Dr. Zach Barth, for critical feedback and thoughtful discussion regarding this project. Thanks to the Glaunsinger Lab for the plasmid encoding 6xHis-SENP2. Special thanks to Denise Schichnes and the UC Berkeley Biological Imaging Facility for advice and equipment. We are incredibly grateful for the Biology Scholars Program (Research Fellows Program) and the McNair Scholars Program for supporting MHTN as an undergraduate researcher, allowing them to pursue scientific research. MHTN extends special gratitude to Ernesto Bonilla and Bear for all their support.

