## Supplemental Figures and Tables for "A phage weaponizes a satellite recombinase to subvert viral restriction"

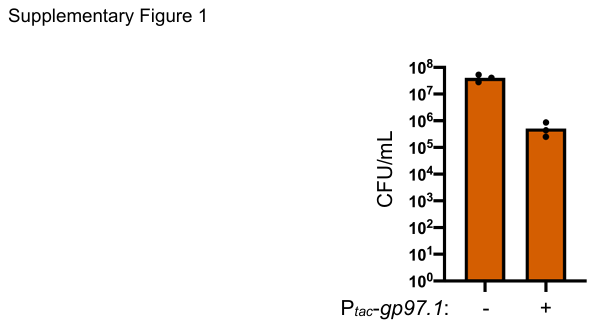

**Figure S1.** *V. cholerae* expressing *gp97.1* exhibited toxicity dependent on the presence of PLE2. Cell viability as measured by colony forming units (CFU) per mL of PLE2(+) *V. cholerae* containing an induced empty vector plasmid control or *gp97.1* (see methods). Bar height represents the mean and dots are measurements from independent assays.

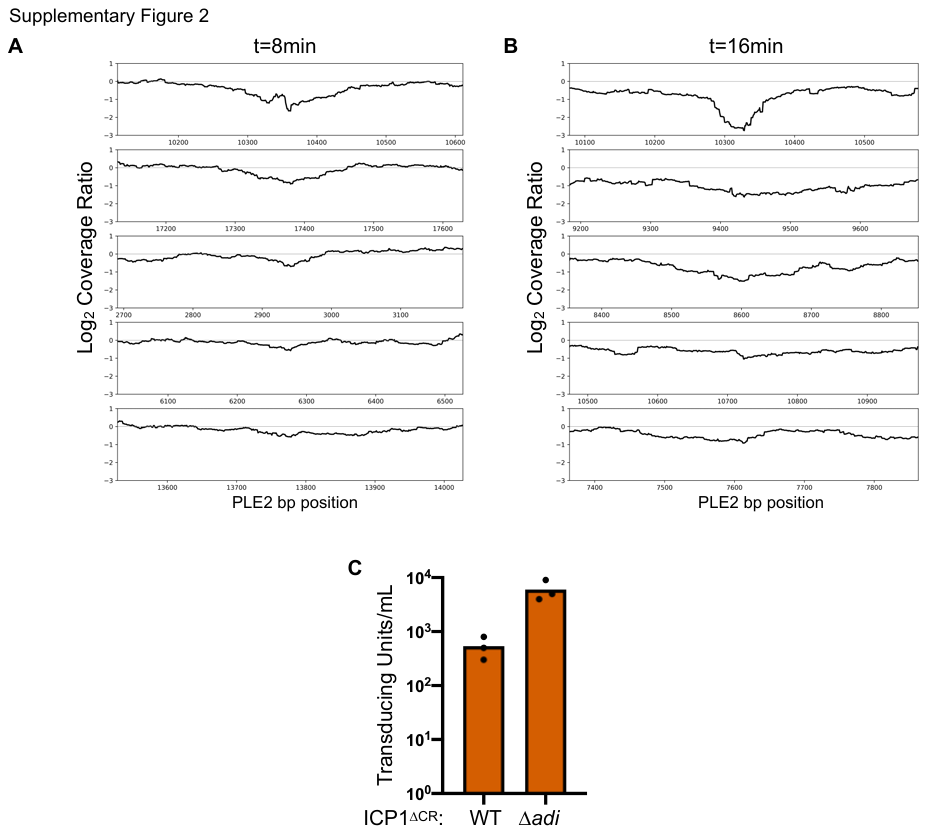

**Figure S2.** (**A, B**) As in (Figure 2B) but showing the five most prominent coverage drops detected at the time indicated following phage infection. The top graph shows the most prominent difference in coverage and is what is shown in Figure 2B. In all panels the peak difference in coverage is centered on the X axis which shows the base pair position in the PLE2 genome. (**C**) Transduction of PLE2 following infection with ICP1 or the ∆*adi* derivative. Bar height represents the mean of three biological replicates and each dot is a measurement from an independent assay.

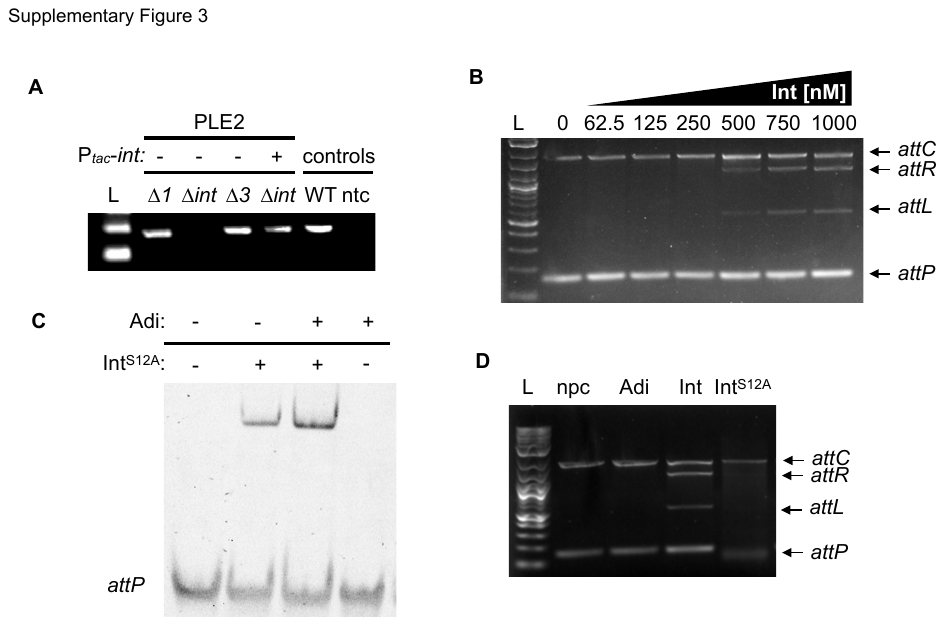

**Figure S3.** (**A**) Agarose gel showing the circularization PCR of PLE2 derivatives lacking ORFs and complemented with a plasmid expressing *int* as indicated, following infection by ICP1^2006∆CR^ (ORFs 1 and 3 flank the integrase and were also tested). Controls on the right show circularization of wild-type PLE2 following ICP1 infection (WT) and a no template control (ntc) for the PCR. (**B**) Agarose gel showing products from *in vitro* DNA recombination assays of *attC* and *attP* (integration) at increasing Int concentrations (as indicated above the gel). The input templates and recombination products (*attL* and *attR*) are labeled on the right. (**C**) Electrophoretic mobility shift assay (EMSA) using a DNA probe containing *attP* incubated with Int^S12A^ and/or Adi. (**D**) Agarose gel showing products from *in vitro* DNA recombination assays of *attC* and *attP* (integration) incubated with no protein (npc), or 500nM of purified Adi, Int, or Int^S12A^. The input templates and recombination products (*attL* and *attR*) are labeled on the right. For (A), (B) and (D), L indicates the ladder.

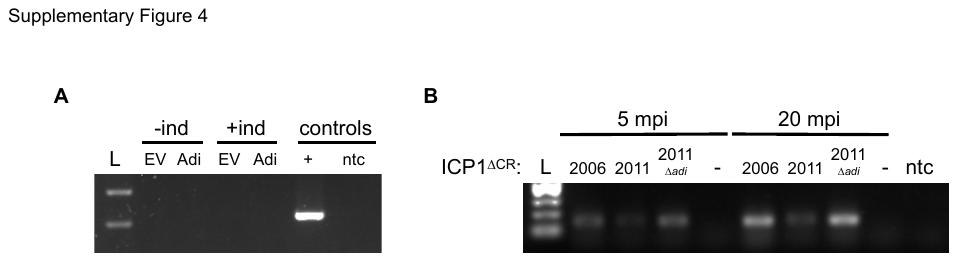

**Figure S4.** (**A**) Agarose gel showing circularization PCR of miniPLE2 (construct containing PLE2 *int* under its endogenous promoter and integrated in the same attachment site as PLE2) with an empty vector (EV) or plasmid expressing *adi* uninduced (-ind) or induced (+ind). Controls on the right include circularization of miniPLE2 following ICP1 infection (+) and a no template control (ntc) for the PCR. (**B**) Agarose gel showing circularization PCR of PLE2 5- and 20-minutes post-infection (mpi) with the ∆CRISPR (CR) derivative of the ICP1 isolate indicated. An uninfected strain with PLE2 was included as a negative control (-) at both time points, and a no template control (ntc) for the PCR was also included.

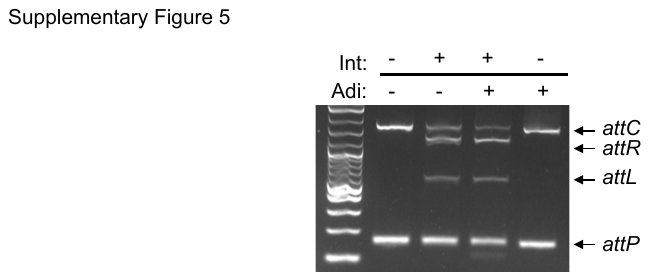

**Figure S5.** Agarose gel of *in vitro* DNA recombination assay of *attP* and *attC* incubated with 500nM of purified Int and/or Adi. The input templates and recombination products (*attL* and *attR*) are labeled on the right.

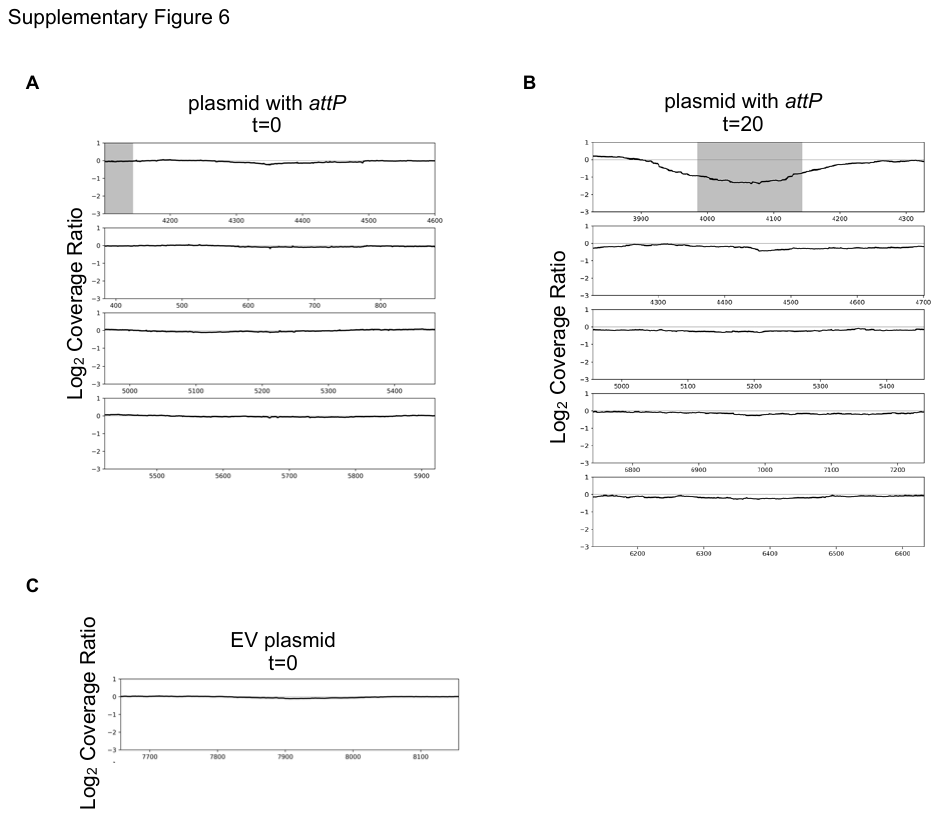

**Figure S6.** As in (Figure 7B) but showing the five most prominent coverage drops detected for the plasmid with *attP* target sequence at t=0 minutes (uninduced), t=20 minutes (induced), and empty vector control at t=0 minutes (uninduced). In all panels the peak difference in coverage is centered on the X axis which shows the base pair position in plasmid. (**A**) Most prominent coverage drops for plasmid with *attP*, only four peaks were detected and are shown. (**B**) Most prominent coverage drops for plasmid with *attP* at t=20 minutes (induced). The top graph shows the most prominent difference in coverage and is what is shown in Figure 7B. The region comprising the circularization junction, *attP*, is highlighted in grey. (**C**) The most prominent coverage drop (and the only one detected) for the empty vector control plasmid at t=0 (uninduced). No panels are shown for the empty vector control plasmid at t=20 minutes (induced) as no peaks were detected.

**Supplementary Table S1: Strains used in this study**

| Strain | Description* | Source |
| --- | --- | --- |
| KDS6 | *V. cholerae* O1, El Tor biotype | Lab collection |
| KDS36 | *V. cholerae* E7946 PLE1 | (13) |
| KS1118 | *V. cholerae* E7946 PLE1::SpecR | (cite) |
| KDS37 | *V. cholerae* E7946 PLE2 | (13) |
| KDS104 | *V. cholerae* E7946 PLE2::kanR | (13) |
| KDS228 | *V. cholerae* E7946 *ΔlacZ*::specR | This study |
| MN35 | *V. cholerae* E7946 PLE2∆*1*::*spec-frt* | This study |
| MN36 | *V. cholerae* E7946 PLE2∆*int*::*spec-frt* | This study |
| KS2076 | *V. cholerae* E7946 PLE2∆*int*::*spec-frt* pKL06::EV | This study |
| KS2077 | *V. cholerae* E7946 PLE2∆*int*::*spec-frt* pKL06.2::PLE2*int* | This study |
| MN37 | *V. cholerae* E7946 PLE2∆*3*::*spec-frt* | This study |
| MN818 | *V. cholerae* E7946 PLE2∆*int*::*spec-frt*-PLE2*attP* pKL06::EV | This study |
| MN811 | *V. cholerae* E7946 PLE2∆*int*::*spec-frt*-PLE2*attP* pKL06.2::PLE2*int* | This study |
| MN812 | *V. cholerae* E7946 PLE2∆*int*::*spec-frt*-PLE2*attP* pKL06.2::PLE2*intS12A* | This study |
| MN573/MN574 | *V. cholerae* E7946 PLE2 pKL06::EV | This study |
| KS2261/KS2262 | *V. cholerae* E7946 PLE2 pKL06.2::ICP1^2011^*gp97.1* | This study |
| KS2096 | *V. cholerae* E7946 PLE1 pKL06.2::PLE2*int* | This study |
| KS2086 | *V. cholerae* E7946 PLE1::PLE2*attP* | This study |
| KS2096 | *V. cholerae* E7946 PLE1 pKL06.2::PLE2*int* | This study |
| KS2097 | *V. cholerae* E7946 PLE1::PLE2*attP* pKL06.2::PLE2*int* | This study |
| MN456 | *V. cholerae* E7946 PLE1::PLE2*attP* pKL06.2::PLE2*intS12A* | This study |
| MN744 | *V. cholerae* E7946 *∆lacZ*::PLE2*attP*-*gfp* pKL06::EV | This study |
| MN745 | *V. cholerae* E7946 *∆lacZ*::PLE2*attP*-*gfp* pKL06.2::PLE2*int* | This study |
| MN746 | *V. cholerae* E7946 *∆lacZ*::PLE2*attP*-*gfp* pKL06.2::PLE2*intS12A* | This study |
| MN640 | *V. cholerae* E7946 ∆*lacZ*::PLE2*int* | This study |
| MN350 | *V. cholerae* E7946 ∆*lacZ*::ICP1^2011^*gp97.1* | This study |
| MN657 | *V. cholerae* E7946 ∆*lacZ*::PLE2*int VCA0692*:: ICP1^2011^*gp97.1* | This study |
| MN744/MN747 | *V. cholerae* E7946 ∆*lacZ*::PLE2*attP* pKL06.2::EV | This study |
| MN745/MN748 | *V. cholerae* E7946 ∆*lacZ*::PLE2*attP* pKL06.2::PLE2*int* | This study |
| MN746/MN749 | *V. cholerae* E7946 ∆*lacZ*::PLE2*attP* pKL06.2::PLE2*intS12A* | This study |
| MN819/MN820 | *V. cholerae* E7946 ∆*lacZ*::PLE2*attP chr::PLE1-SpecR* pKL06.2::EV | This study |
| MN821/MN822 | *V. cholerae* E7946 ∆*lacZ*::PLE2*attP chr::PLE1-SpecR* pKL06.2::PLE2*int* | This study |
| MN395/MN398 | *V. cholerae* E7946 ∆*lacZ*:: ICP1^2011^*gp97.1* pKL06::EV | This study |
| MN396/MN399 | *V. cholerae* E7946 ∆*lacZ*:: ICP1^2011^*gp97.1* pKL06.2::PLE2*int* | This study |
| MN397/MN400 | *V. cholerae* E7946 ∆*lacZ*:: ICP1^2011^*gp97.1* pKL06.2::PLE2*intS12A* | This study |
| MN805 | *V. cholerae* E7946 ∆*lacZ*::PLE2*intS12A VCA0692*:: ICP1^2011^*gp97.1 ∆attC::SpecR* pKL06.2::EV | This study |
| MN806 | *V. cholerae* E7946 ∆*lacZ*::PLE2*intS12A VCA0692*:: ICP1^2011^*gp97.1 ∆attC::SpecR* pKL06.2::PLE2*attP* | This study |
| MN807 | *V. cholerae* E7946 ∆*lacZ*::PLE2*int VCA0692*:: ICP1^2011^*gp97.1 ∆attC::SpecR* pKL06.2::EV | This study |
| MN808 | *V. cholerae* E7946 ∆*lacZ*::PLE2*int VCA0692*:: ICP1^2011^*gp97.1 ∆attC::SpecR* pKL06.2::PLE2*attP* | This study |
| MN116/MN117 | *E. coli* BL21 ICP1^2011^*gp97.1*-*pSUMO* | This study |
| MN118/MN119 | *E. coli* BL21 PLE2*int*-*pSUMO* | This study |
| MN120 | *E. coli* BL21 PLE2*intS12A*-*pSUMO* | This study |
| MN392 | *E. coli* BL21 6xHis-SENP2 | This study |
| KSphi39 | ICP1_2011_A | (18) |
| ACMphi232 | ICP1_2011_A ∆CRISPR ∆spacer2-9 *cas1*^D244A^ | (19) |
| KSphi117 | ICP1_2011_A ∆CRISPR ∆spacer2-9 *cas1*^D244A^ ∆*gp97.1* | This study |
| KSphi38 | ICP1_2006_E ΔCRISPR Δ*cas2-3* | (13) |
| ACMphi289 | ICP1^1992^ | (23) |
| MNphi10-13 | ICP1^1992^ ∆*gp97.1* | This study |
| ACMphi258 | ICP1_2017_D ∆*cas2-3* | (19) |
| KSphi161 | ICP1_2019_Mat_C ∆*cas2-3* | (23) |
| MNphi14-17 | ICP1_2019_Mat_C ∆*cas2-3* ∆*gp97.1* | This study |

**Supplementary Table S2: Primers used in this study**

| Primer | Sequence | Purpose | Source |
| --- | --- | --- | --- |
| zac68 | CTGAATCGCCCTACCCGTAC | qPCR PLE | (12) |
| zac69 | GTGAACCAACCTTTGTCGCC | qPCR PLE | (12) |
| MN262 | GATAGGTCATTTTATACTTCTGACAAAGTTACATATGACTTGCATTAGTG | F sequence EMSA probe | This study |
| MN263 | CACTAATGCAAGTCATATGTAACTTTGTCAGAAGTATAAAATGACCTATC | R sequence EMSA probe | This study |
| MN69 | TAACTACTGTTTGTACCAG | F primer to amplify *attP/attR* probe for *in vitro* recombination | This study |
| MN70 | AATTTAAGATCCTTCTAACATAGTG | R primer to amplify *attP/attL* probe for *in vitro* recombination | This study |
| MN168 | AACGTATTTTTCGATGTGGC | F primer to amplify *attC/attL* probe for *in vitro* recombination | This study |
| MN169 | ATCGAAATAGAAGCGATTGG | R primer to amplify *attC*/*attR* probe for *in vitro* recombination | This study |

**Supplementary Table S3: Sequence of PLE2*attP***

| Sequence | Purpose | Source |
| --- | --- | --- |
| ACCAGATTAACTACTAGATTAATGCCATATATGAGCCGCTACCTTTCGAGGTGGCGGTTTTTGTTTTGTGTAATCTCAAAAGTTACATATGACTTGCATTAGTGGATAGGTCATTTTATACTTCTGACAAATCACTATGTTAGAAGGATCTTAAATTATG | Cloned into GFP reporter, PLE1, and plasmid as target sequence | This study |
